# Systematic analysis of RhoGAP expression and function in border cell morphology and migration

**DOI:** 10.64898/2026.04.07.717016

**Authors:** Abhinava K. Mishra, Emily G. Gemmill, Joseph P. Campanale, James A. Mondo, Veronique Lisi, Kenneth S. Kosik, Denise J. Montell

## Abstract

Rho family GTPases are central hubs in the signaling and cytoskeletal networks that govern cell morphology and behavior. GTPase-activating proteins (GAPs) inactivate them by accelerating GTP hydrolysis. However, a systematic analysis of GAPs in cell migration is lacking. Here, we report screens for RhoGAP expression and function in migratory Drosophila border cells. Constitutively active Cdc42, Rac, or Rho causes defects, demonstrating that negative regulation is critical. Integrating single-cell RNAseq with published datasets reveals that most of the 22 RhoGAPs are expressed in border cells. RNAi knockdown shows most RhoGAPs are functionally required. We developed automated image analysis tools to sensitively and objectively classify border cell morphologies, defining a normal morphological phase space. RhoGAP perturbations push clusters outside this range. In-depth analysis of RhoGAPp190 reveals that loss-of-function resembles Rho hyperactivation and gain-of-function resembles myosin II inhibition. Thus, complex spatiotemporal sculpting of RhoGTPase activities requires diverse RhoGAPs within a single cell type to control morphology and motility *in vivo*.

## Introduction

Collective cell migration drives embryonic morphogenesis, tissue repair, and cancer invasion (Lintz et al., 2017; Scarpa and Mayor, 2016; Friedl and Mayor, 2017; Rørth, 2009; Mayor and Etienne-Manneville, 2016; Campanale and Montell, 2023; Te Boekhorst et al., 2016). During migration, Rho family GTPases are central nodes in the signaling networks that control and coordinate polarity, adhesion, and actin dynamics within and between moving cells (Zegers and Friedl, 2014; Fukata et al., 2003; Ridley, 2001; Raftopoulou and Hall, 2004; Lawson and Ridley, 2018; Lawson and Burridge, 2014). Rho, Rac, and Cdc42 cycle between inactive, GDP-bound, and active, GTP-bound states. Guanine nucleotide exchange factors (GEFs) promote activation, whereas GTPase-activating proteins (GAPs) accelerate GTP hydrolysis to terminate signaling.

In mammals, more than 60 RhoGAPs and ∼80 RhoGEFs greatly outnumber the ∼20 widely-expressed Rho GTPases, suggesting that regulation of GTPase activity is complex (Mosaddeghzadeh and Ahmadian, 2021). However, the nature of that complexity is poorly understood (Dahmene et al., 2020; Hodge and Ridley, 2016). Do cells express all these regulators simultaneously, or do different cells express distinct subsets? Do GAPs function redundantly or contribute specialized functions? Negative regulation of GTPase activities via RhoGAPs is likely critical for cell morphology and motility (Bidaud-Meynard et al., 2019). Yet, only a minority of RhoGAPs have been functionally characterized.

Drosophila provides a tractable model in which to analyze RhoGAP function *in vivo*. The fly genome encodes five canonical Rho GTPases, 26 GEFs, and 22 GAPs. Several Drosophila RhoGAPs have roles in morphogenesis and tissue dynamics (Mason et al., 2016; Jackson et al., 2024) (Fic et al., 2020) (Billuart et al., 2001). A systematic analysis of RhoGAP expression during epithelial cell division has revealed some dynamic subcellular localizations (di Pietro et al., 2023), but studies of migratory cells are lacking.

Border cells have been extensively used to study Rho-family GTPase function in collective migration (Zegers and Friedl, 2014; Wu et al., 2011; Montell, 2003; Montell et al., 2012; Rørth, 2009) and notably provided the first demonstration of the role of Rac in cell migration (Murphy and Montell, 1996). During Drosophila oogenesis, border cell clusters consist of 6-8 follicle cells that migrate within a structure called an egg chamber, from the anterior pole, between nurse cells, to the oocyte (Montell et al., 1992)**(Fig. 1A)**. These clusters comprise 4-6 migratory cells surrounding two non-motile polar cells. Leader cells extend protrusions in the direction of migration (Murphy and Montell, 1996) while follower cells crawl on one another and on nurse cells to propel the cluster forward (Campanale et al., 2022). Rac1, Rac2, and Mtl function redundantly to regulate border cell protrusion and migration (Murphy and Montell, 1996; Geisbrecht and Montell, 2004), downstream of RTK guidance receptors and multiple GEFs (Fernández-Espartero et al., 2013; Duchek et al., 2001). Rac activity is higher in leader cells extending protrusions (Wang et al., 2010; Mishra et al., 2019; Inaki et al., 2012) but is present and required in all migratory border cells (Campanale et al., 2022; Montell et al., 2012; Campanale and Montell, 2023). Rho1 regulates cluster shape and motility (Mishra et al., 2019; Bastock and Strutt, 2007). Cdc42 regulates cell polarity and coordinates cell-cell communication within the cluster (Colombié et al., 2017). Despite this understanding of positive regulation by GEFs, negative regulation by RhoGAPs remains poorly understood. RhoGAP18B modulates F-actin organization and migration in border cells (Lei et al., 2023). Constitutively active Rho, Rac, or Cdc42 produce distinctive defects, demonstrating that negative regulation is critical; however, there has been no systematic analysis of which RhoGAPs are expressed or required in this collective, nor how they contribute to distinct aspects of morphology and motility.

**Figure 1:**
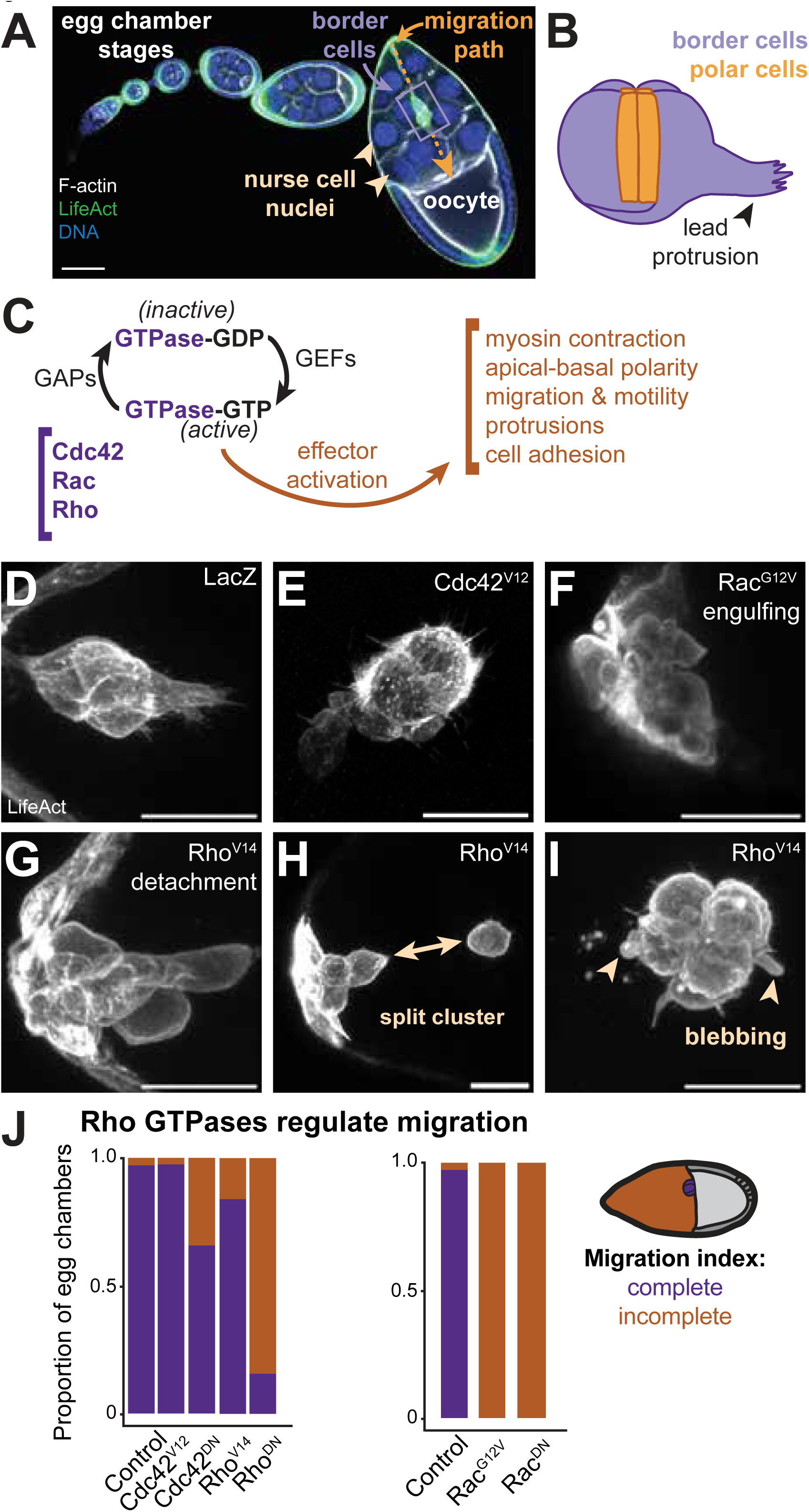
Negative regulation of Rho GTPases is essential for border cell morphology and migration. (A) Confocal micrograph of an ovariole labeled with phalloidin (white) to stain F-actin, Hoeschst (Blue, DNA), and LifeActinGFP driven by c306Gal4. The border cell cluster (purple box), migration path (orange arrow), and nurse cell nuclei (yellow arrows) are indicated. Scale bar = 50μm **(B)** Border cell clusters consist of 4-6 outer, migratory cells and 2 central, nonmotile polar cells **(C)** Rho GTPase activity cycle regulated by GEFs and GAPs. **(D-I)** Confocal images of representative border cell clusters expressing LifeActGFP directly driven by the ovarian enhancer from the *slbo* gene (*slbo*LifeActGFP, white) together with *slbo*Gal4 and the indicated UAS-transgenes. **(D)** Control cluster initiating migration. **(E)** Cluster expressing Cdc42^V12^ **(F)** Rac^V12^, **(G-I)** Rho^V14^, **(J)** Migration defects caused by inhibition or hyperactivation of Cdc42, Rho, or Rac. DN: dominant-negative. Scale bars = 20μm.

Here, we systematically analyze RhoGAP expression and function in border cells. We combine single-cell RNAseq with published datasets to show that most RhoGAPs are expressed in border and/or polar cells. RNAi-mediated knockdown reveals that most contribute to normal morphology and migration. To objectively quantify morphological phenotypes, we develop automated image analysis tools measuring circularity, protrusiveness, and protrusion directionality. This reveals that wild-type border cells adopt shapes within a defined morphological phase space and that different GAP knockdowns push clusters outside this range in distinct ways, suggesting specialized functions. Analysis of RhoGAPp190 reveals phenotypes resembling Rho hyperactivation upon knockdown and myosin II inhibition upon overexpression, consistent with p190 negatively regulating Rho-ROCK-myosin signaling. Overexpression of p190 produces extreme morphologies due to loss of cell-cell adhesion within the collective. These results demonstrate that complex spatiotemporal sculpting of RhoGTPase activities requires multifarious RhoGAPs acting within a single cell type. Our systematic approach provides a comprehensive catalog of functional requirements and a methodological framework for dissecting RhoGAP complexity in collective migration.

## Results

### Negative regulation of RhoGTPases is essential for border cell morphology and migration

Drosophila oocytes grow and develop through 14 stages within egg chambers composed of 16 germline cells (15 nurse cells and 1 oocyte), surrounded by a monolayer of ∼800 somatic epithelial follicle cells (**Fig. 1A**) (King, 1970). Border cells are a subset of follicle cells that migrate collectively during stage 9, from the anterior pole, in between nurse cells, to the oocyte (**Fig. 1A**), where they attach by stage 10. These clusters are composed of 4-6 migratory cells that surround and carry two non-motile polar cells (**Fig. 1B**). One or two leader cells explore their surroundings by extending and retracting protrusions predominantly in the direction of migration (Murphy and Montell, 1996; Fulga and Rørth, 2002; Dai et al., 2020). Three-five follower cells crawl on one another and the substrate nurse cells to propel the cluster forward (Campanale et al., 2022). Thus, wild-type clusters dynamically remodel the cytoskeleton to change shape as they move. Cdc42, Rac1, Rac2, Mtl, and Rho all contribute to normal cluster morphology and/or motility (**Fig. 1C**).

Hyperactivity of Rho family GTPases, especially Rac and Rho, also perturbs border cell cluster morphology and motility. Whereas control border cell clusters are typically compact and well-organized with a single major protrusion as they initiate migration **(Fig. 1B, D)**, border cells expressing hyperactive GTPases exhibit altered morphologies. We used *slbo*Gal4 (Rørth et al., 1998) to drive expression of UAS-Cdc42^V12^, UAS-Rac1^V12^, or UAS-Rho^V14^ in outer, migratory border cells just prior to migration. Hyperactive Cdc42 (Cdc42^V12^) caused the mildest phenotype with relatively normal cluster cohesion **(Fig. 1E)** and migration to the oocyte (Murphy and Montell, 1996). Rac^V12^ expression **(Fig. 1F)** blocks most protrusions, severely impairs movement (Geisbrecht and Montell, 2004), and renders border cells phagocytic toward polar cells and nurse cells (Mishra et al., 2023). Rho^V14^-expressing border cells are typically rounder than normal **(Fig. 1G-I)** and less adherent to one another (Bastock and Strutt, 2007; Aranjuez et al., 2016; Gabbert et al., 2023), to the point where clusters sometimes split apart **(Fig. 1H)**. Blebs also frequently form **(Fig. 1I)**. Despite the dramatic changes in cell and cluster morphology, Rho^V14^-expressing clusters exhibit surprisingly mild migration defects **(Fig. 1J)**, presumably due to switching from an adhesive, mesenchymal migration mode to a contractility-driven motility (Friedl and Wolf, 2010; Mishra et al., 2019). Cdc42^V12^-expressing clusters complete migration normally, whereas clusters expressing either dominant-negative or constitutively active Rac exhibit the most severe migration defects **(Fig. 1J)**.

### Most RhoGAPs are expressed in border and/or polar cells

While RhoGTPase activity requires complex spatiotemporal regulation during collective cell migration generally and border cell cluster migration specifically, how many of the >20 RhoGAP genes are expressed in border cells was unknown. We carried out a single-cell RNA sequencing (scRNAseq) analysis on hand-dissected, egg chambers that were late stage9/early stage 10 **(Fig. 2A)** expressing nuclear dsRed specifically in polar cells **(Fig. 2A, B)** and LifeActGFP in border cells (**Fig. 2B’**), centripetal cells, and posterior follicle cells **(Fig. 2A)**. Egg chambers were dissociated into single cells and subjected to DropSeq (Macosko et al., 2015)**(Fig. S2A-C)**. Unsupervised clustering revealed a variety of cell populations (**Fig. 2C**). We then used dsRed expression to identify polar cells unambiguously. 88% of the dsRed-positive cells were located within cluster 21 **(Fig. 2C and Fig. S2)**. We then used GFP expression as well as *slbo* and *fruitless,* which are mRNAs known to be enriched in border cells, to identify a candidate border cell population (**Fig. 2D**). We found that a majority (58%) of the cells expressing GFP, *fruitless*, and *slbo* populated cluster 7 **(Fig. 2D)**. These results are consistent with the fact that polar cells differentiate from other follicle cells early in oogenesis whereas border cells differentiate at stages 8/9.

**Figure 2:**
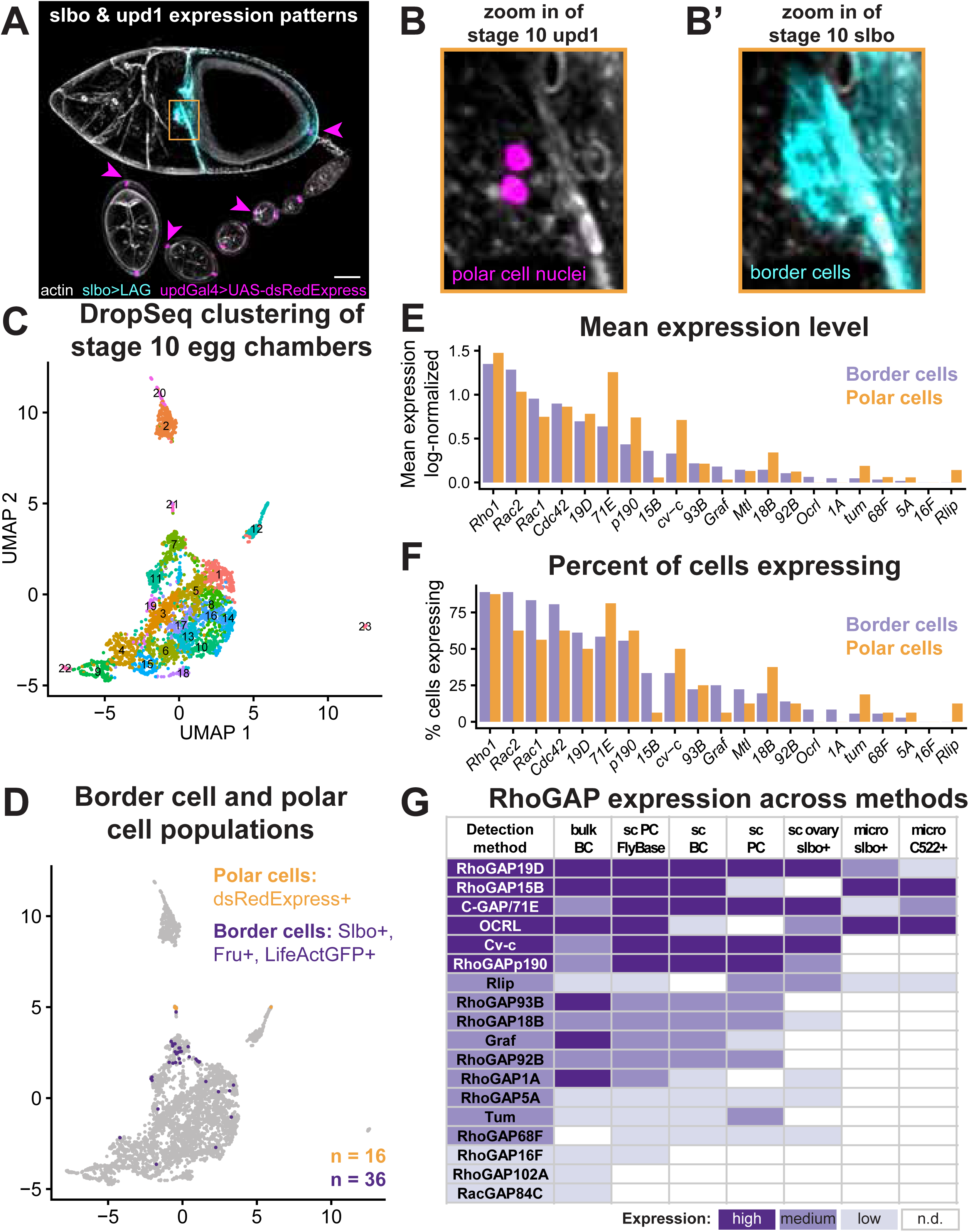
Most RhoGAPs are expressed in border cell and/or polar cells. (A) *Slbo* and *upd1* expression patterns **(B)** Enlargements of panel A showing the **(B’)** polar cells and **(B’’)** border cells **(C)** Clustering of stage 10 egg chambers shows a variety of cell populations, but none fully and exclusively capture border or polar cells. **(D)** Border cells are selected for using slbo-, fruitless-, and LifeActGFP-positive cells, and polar cells are determined using dsRedExpress-positive cells. **(E)** Mean expression of GTPases and GAPs in the border and polar cell populations. Gene expression values were log-normalized to account for differences in sequence depth between cells. Values represent log(count/total counts per cell x 10,000). **(F)** Percent expression of GTPases and GAPs in the border and polar cell populations. **(G)** RhoGAP mRNA expression shows that RhoGAPs are expressed at different levels and that more highly expressed GAPs are detected in more experiments. High, medium, and low bins were defined as relative levels of expression within each experiment. n.d. = not detected

We then analyzed the expression of Rho GTPases and RhoGAPs in the 36 border cells and 16 polar cells that passed quality control as well as in all cells in clusters 7 and 21 **(Fig. 2D and Fig. S2F and S2G)**. Rho1, Rac1 and Rac2, and Cdc42 were relatively highly expressed in both border and polar cells, followed closely by RhoGAP-19D, -71E, -p190, -15B, and Cv-c (**Fig. 2E and 2F**). There were some differences between polar cells and border cells; for example, RhoGAP71E was more highly expressed in polar cells. However, overall, the expression patterns were similar. The mean expression per cell (**Fig. 2E)** correlated exceptionally well with the percent of cells expressing each gene (**Fig. 2F and Fig. S2H,** R^2^>0.96).

Several previously published studies have reported gene expression analyses of border and/or polar cells, each with a different experimental design. Consistent with our results, unsupervised clustering identified polar cells more effectively than border cells. Flybase reports scRNA seq for polar cells but not border cells. Rust et al (Rust et al., 2020) reported scRNAseq on whole ovaries, within which they identified a putative border cell population based on *slbo* expression. Burghardt et al sorted *slbo*Gal4-expressing cells from early, mid, and late stage 9 egg chambers and carried out bulk RNAseq rather than single-cell (Burghardt et al., 2023). Earlier studies used microarray analysis to evaluate border and/or polar cell gene expression profiles (Wang et al., 2006; Borghese et al., 2006). Therefore, we compared RhoGAP expression across these datasets. Expression of each of the RhoGAP genes was detected in border and/or polar cells by at least one method **(Fig. 2G)**. Furthermore, more abundant RhoGAP mRNAs were detected in more experiments. This pattern argues that the absence of detection by any single method does not indicate the absence of expression; rather, the integration of multiple complementary approaches provides the most comprehensive view of the RhoGAP expression landscape **(Fig. 2G)**.

As expected, the two microarray analyses, while well-correlated, were less sensitive and therefore detected expression of fewer RhoGAP genes than RNAseq experiments. Bulk RNAseq of isolated stage 9 *slbo*Gal4-expressing cells was slightly more sensitive than the scRNAseq methods. While our scRNAseq yielded a relatively modest number of border and polar cells passing quality control, this small sample size proved sufficient for detecting gene expression patterns in rare cell populations. Furthermore, our findings that most RhoGAP mRNAs are expressed in border and/or polar cells validate and are validated by the published, independent gene expression analyses **(Fig. 2G)**. We conclude that most or all RhoGAP genes are expressed in border and/or polar cells, and the expression levels vary.

### Many RhoGAP RNAi knockdowns caused migration defects

Since most or all of the RhoGAP mRNAs were expressed at some level in border and/or polar cells, we systematically analyzed the effects of inhibiting RhoGAP expression on border cell migration (**Fig. 3**). We used c306Gal4, which is expressed in outer, migratory border cells and polar cells (**Fig. 1A**) to express multiple UAS-RNAi lines against each gene, when available. c306Gal4 is also expressed earlier in development than slboGal4, allowing more time for the RNAi to be effective. Crosses were grown at 18 ℃, and all border cells expressed *slbo*LifeActGFP. Female flies were fattened for 72 hours at 30 ℃ and their ovaries dissected, fixed, and stained for nuclei (Hoechst) and F-actin (Phalloidin). These were then imaged with a confocal microscope to quantify migration defects at stage 10 (**Fig. 3A**). Representative images are shown (**Fig. 3B**). In keeping with the expression data, migration defects were observed for many lines (**Fig. 3C**). Some RNAi lines produced stronger phenotypes than others even when targeting the same gene, demonstrating the importance of testing as many lines as possible to achieve the greatest sensitivity.

**Figure 3:**
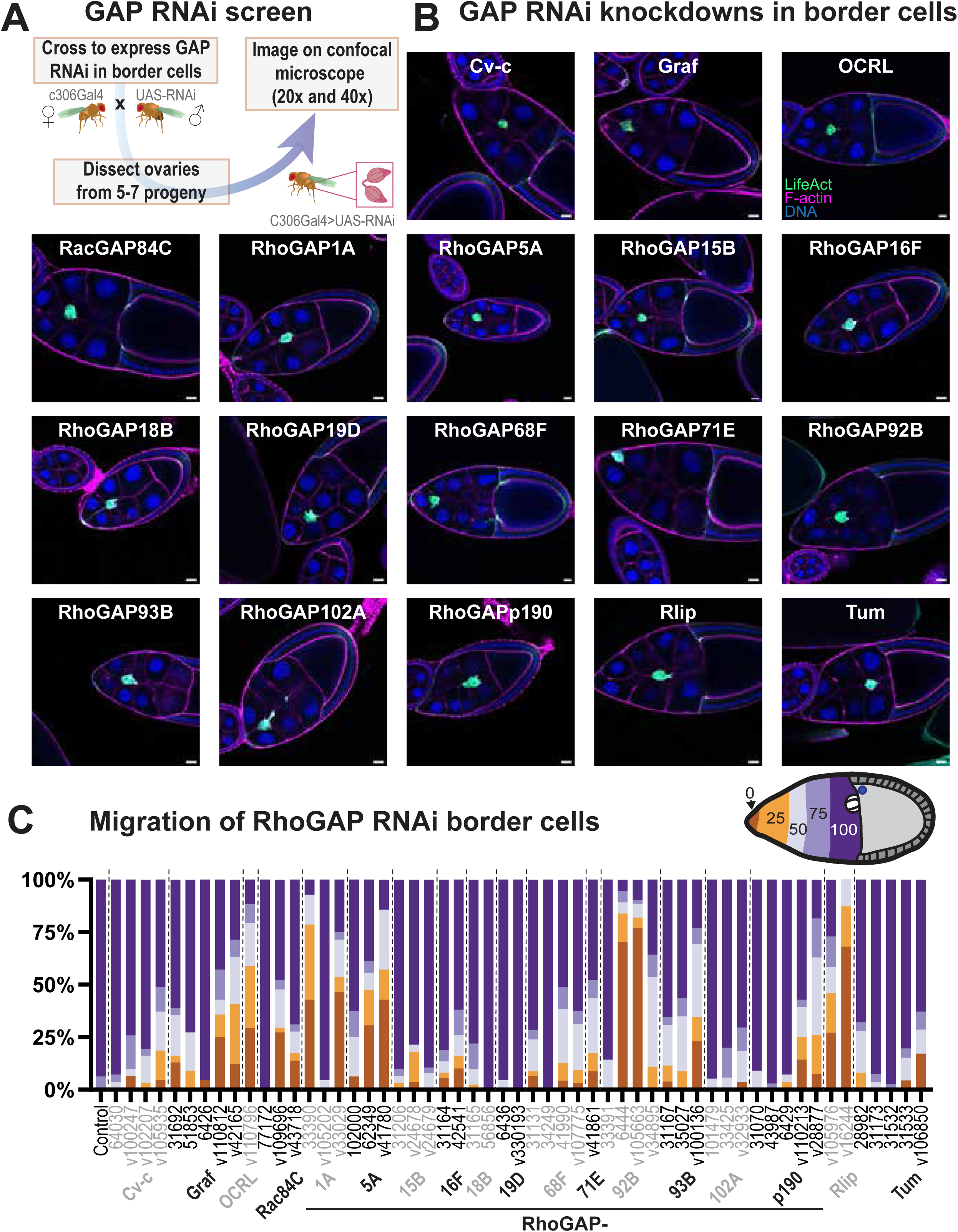
Many RhoGAP RNAi knockdowns caused migration defects. (A) Workflow for RNAi screen in *Drosophila* border cells. (B) Examples of GAP RNAi knockdowns in border cells (20x). (C) Migration defects quantified at stage 10. The fraction of clusters that reached the oocyte by stage 10 is indicated in dark purple. The fraction that did not detach is indicated in dark brown. Those with milder defects are indicated by lighter colors. The GAP genes are arranged alphabetically from left to right, with multiple RNAi lines for most GAPs. Scale bars = 20 μm.

RNAi-mediated knockdown has well-known caveats, including variable knockdown efficiency and potential off-target effects. To address these concerns, we tested multiple independent RNAi lines for most RhoGAPs. Phenotypes observed with multiple independent lines targeting the same gene provide strong evidence against off-target effects. Conversely, the absence of detectable phenotypes could reflect either a functional redundancy with other GAPs or insufficient knockdown, particularly for highly expressed genes. The observations that most GAPs were both expressed and required support the validity of the two analyses. Interestingly, expression level did not always correlate with phenotypic severity. In fact, RhoGAP19D was highly expressed, but neither RNAi line caused a significant defect. This could either be because RhoGAP19D is not required or because the high level of expression is difficult to deplete effectively by RNAi. Conversely, RacGAP84C was only barely detected and in only one of the RNAseq analyses; yet two RNAi lines caused a moderate migration defect. RhoGAP92B was detected at a relatively low level in four RNAseq experiments, and two different RNAi lines cause similarly strong migration defects. The 84C and 92B results could either represent off-target RNAi effects or that RNAi-mediated knockdown is more effective for mRNAs that are expressed at low levels. The latter seems more likely given that two different RNAi lines with no predicted off-targets produced similar border cell migration defects. In other cases, expression level and migration defects correlated better. For example, RhoGAP68F mRNA was detected at a low level in four different expression analyses, and 4/5 RNAi lines caused similarly mild migration defects.

### RhoGAP knockdowns that cause cluster splitting

Some RNAi lines that did not cause strong migration defects nevertheless caused cluster splitting reminiscent of that seen in slboGal4;UAS-Rho^V14^ **(Fig. 1H)**. Whereas control border cells typically migrate as a coherent cluster of cells **(Fig. 4A)**, RhoGAP18B 31165-expressing clusters showed frequent cluster splitting **(Fig. 4B)**. Seven additional RhoGAP RNAi lines also exhibited more frequent cluster splitting than the white RNAi controls **(Fig 4C-F)**.

**Figure 4:**
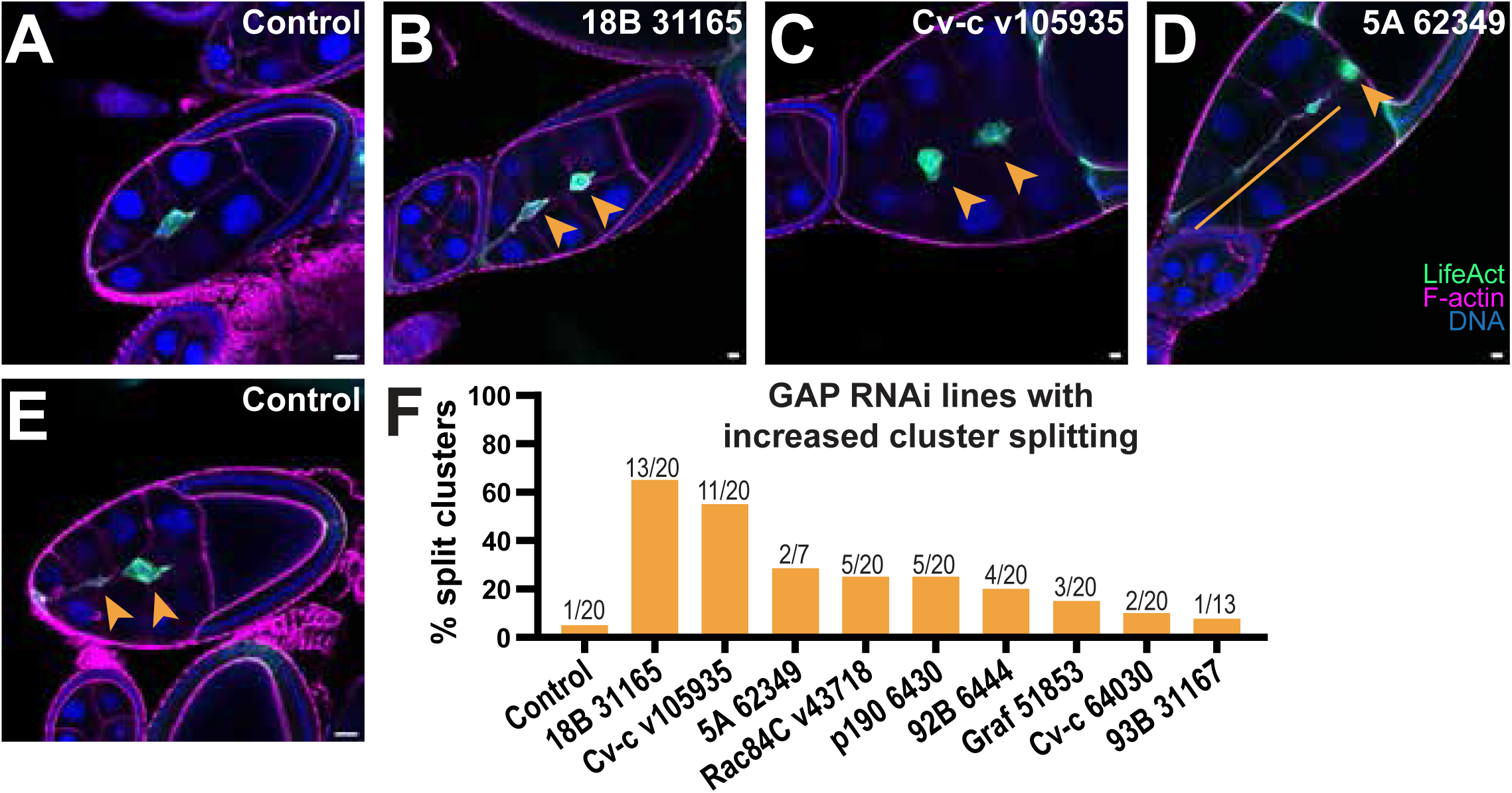
RhoGAP knockdowns that cause cluster splitting. (A) Wildtype clusters typically migrate as a single cluster. (B-F) Cluster splitting (orange arrows) in the indicated genotypes, including a rare example in a control (E). (F) Percentages of border cell clusters that split for the indicated RNAi lines. The number of phenotypic egg chambers/total egg chambers examined is shown above the bars. Scale bar = 20 μm.

### Effects of RhoGAP knockdowns on border cell morphology

We observed a range of morphological phenotypes **(Fig. 5A-S)** that did not always correlate with the severity of migration defects. For example, Rho^V14^ causes dramatic morphological changes—loss of cell-cell adhesion, cluster splitting, and blebbing—yet produces only mild migration defects, likely because cells switch from adhesive to contractility-driven migration modes. To capture morphological phenotypes more sensitively, we developed automated image analysis tools.

**Figure 5:**
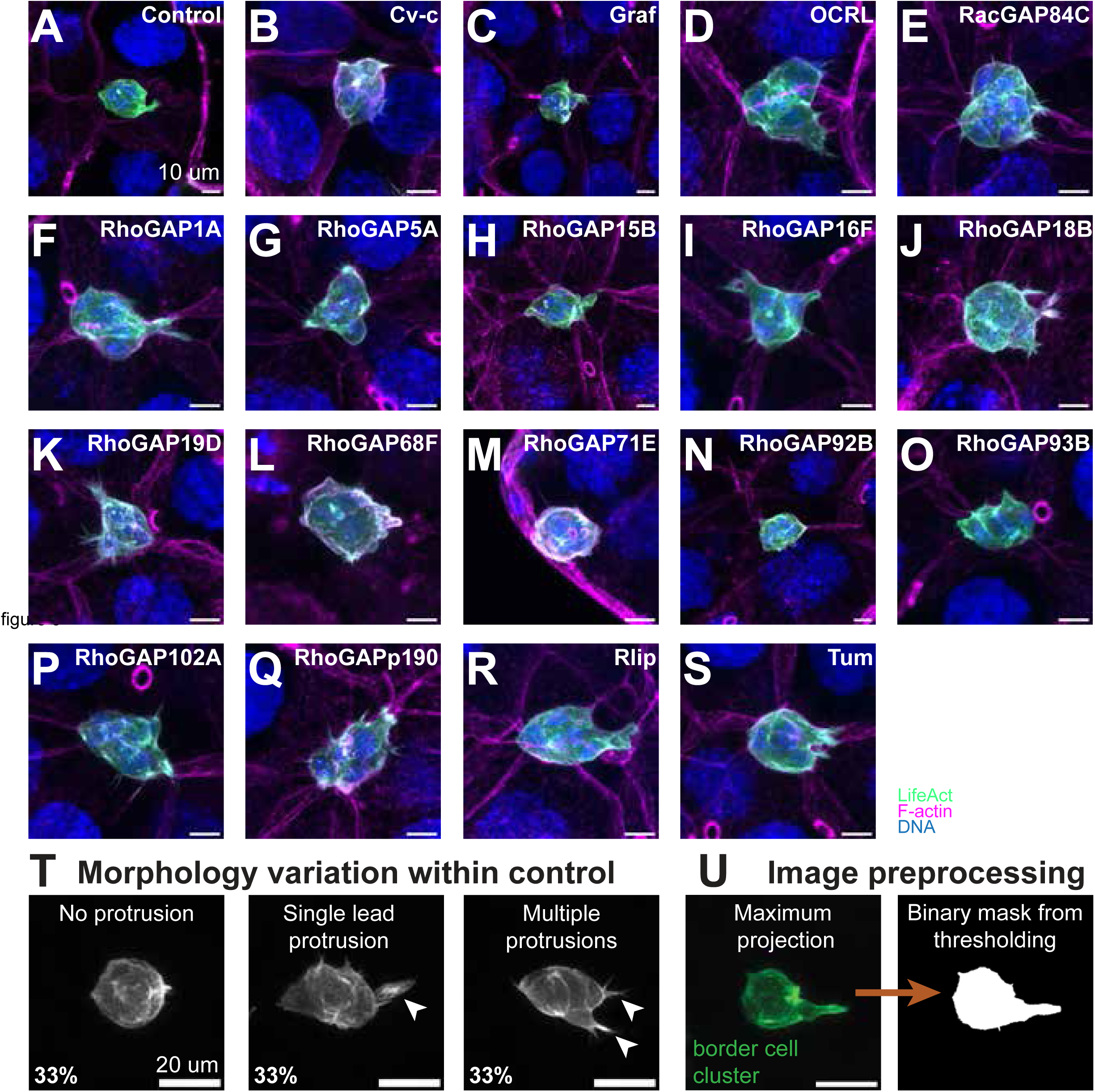
Effects of RhoGAP knockdowns on border cell morphology. (A) A control border cell cluster has one lead protrusion and a rounded cluster body. **(B - S)** Representative photos of GAP knockdowns. Scale bar = 10 μm. **(T)** Within the control, there is a range of protrusions; however, a single lead protrusion is most frequently observed. **(U)** To process these data, the clusters were max projected in FIJI, and then segmented using Li thresholding in Python scikit-image to then allow for downstream analysis. Scale bar = 20 μm.

It was challenging to objectively describe - much less quantify - the varied phenotypes. So, we built a pipeline to describe border cell morphologies objectively and quantitatively and to compare the morphologies to the normal range of shapes that control clusters exhibit as they dynamically extend and retract protrusions (**Fig. 5T**). To this end, we segmented and binarized maximum intensity projections for each replicate across the GAP RNAi lines for further downstream analysis **(Fig. 5U)**.

Importantly, this morphological analysis was restricted to clusters that were migrating, because the shapes of detaching or reattaching clusters are affected by neighboring cells. This criterion necessarily excluded GAP knockdowns that cause the strongest migration defects, as these clusters often fail to complete detachment. Thus, migration and morphology provide complementary readouts: migration defects capture severe functional impairments, while morphological analysis sensitively detects subtler phenotypes in motile clusters.

### Effects of RhoGAP inhibition on border cell protrusion number and directionality

To quantify cluster protrusiveness compared to the control, we projected the segmented clusters onto a circular coordinate system (**Fig. 6A**). Protrusions were defined by any point that extended ≥20% outside of the mean cluster radius (height threshold) and with a prominence threshold >50% (see methods for details). These criteria excluded noise, blebs, broad flat regions, or whole cells that were separated from the cluster. Ectopic protrusions were defined as those for which the tip was located >±20° from the oocyte vector (defined as 0°). All individual protrusions were counted across the control and compiled into a circular histogram that was normalized by the number of replicates to create a composite histogram (**Fig. 6B**).

**Figure 6:**
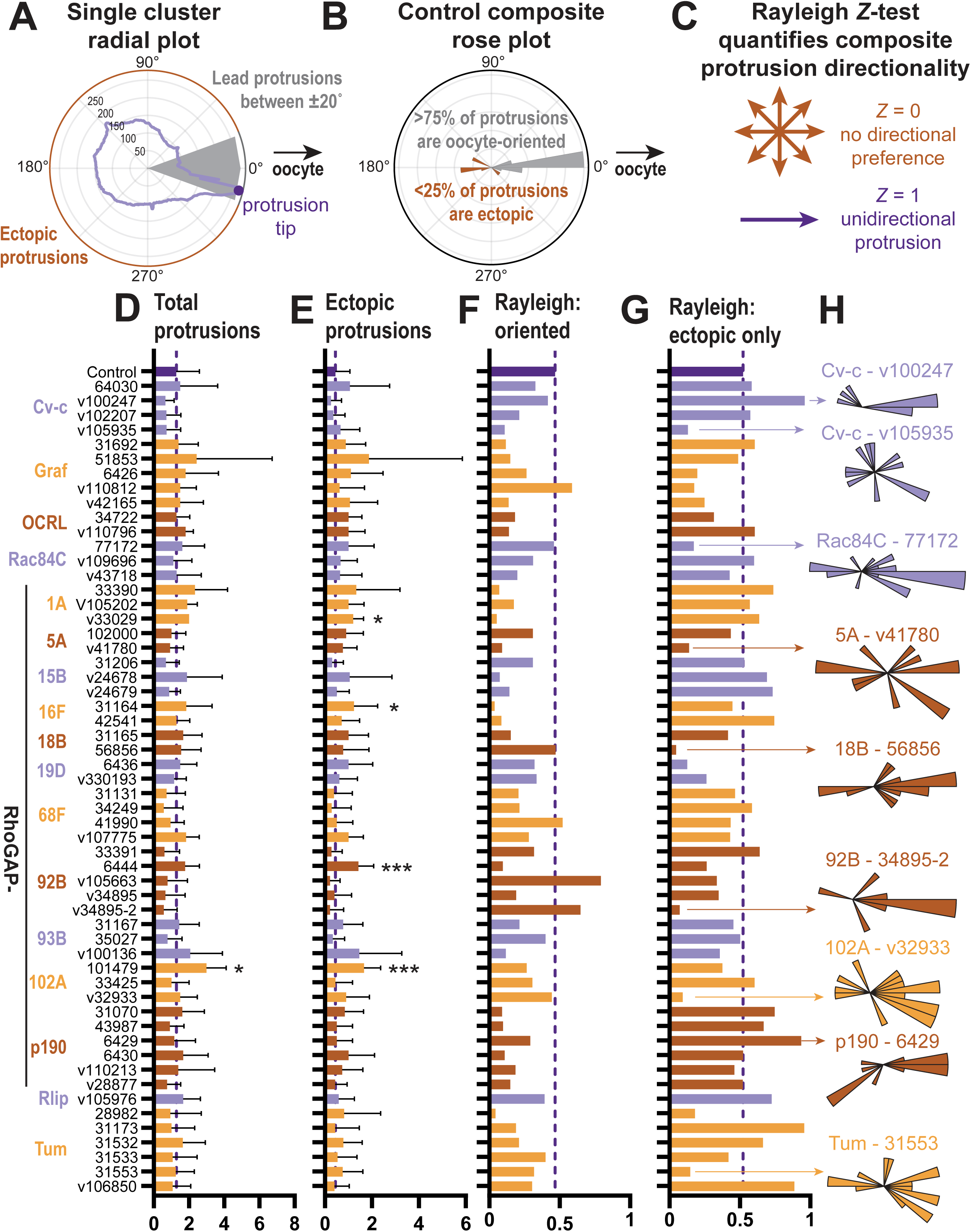
Effects of RhoGAP inhibition on border cell protrusion number and directionality. (A) Individual clusters are projected onto a polar coordinate system (rose plot) to count protrusions. Ectopic protrusions are defined as anything outside of a ±20° region. A protrusion is quantified as anything that protrudes more than 25% past the radius of the cluster and is 50% higher than its nearest neighbor. Units = pixels. **(B)** Histograms on the polar coordinates count the number of protrusions in 10° bins, and these values are normalized by the number of clusters per genotype. **(C)** Rayleigh directionality index measures whether protrusions are unidirectional (*Z* = 1) or uniformly distributed across all angles (*Z* = 0). **(D)** Total protrusions quantified using the rose plot analysis across RhoGAP knockdowns. **(E)** Ectopic protrusions for GAP knockdowns, excluding ±20° from the oocyte vector. **(F)** Rayleigh numbers are low before orienting in the direction of migration. **(G)** After orienting in the direction of migration, clusters exhibit more directional protrusions, and those without directional protrusions are more apparent. **(H)** Rayleigh numbers for ectopic protrusions quantify the directionality of non-functional protrusions. Bars show the mean ± SD. Significance reported using a Bayesian ANOVA. Bayes Factor BF_10_ >3 (*), BF_10_>10 (**), BF_10_ >30 (***).

We used a Rayleigh *Z* test to measure protrusion directionality (**Fig. 6C**). If protrusions are uniformly distributed in all radial directions, *Z* = 0. If protrusions are precisely unidirectional, *Z* = 1. All protrusions were quantified across control and all RNAi lines (**Fig. 6D**). We also separated out ectopic protrusions (**Fig. 6E**). The more strongly oriented the protrusions were toward the oocyte, the higher the *Z* value. Conversely, RNAi lines that caused protrusions to be more randomly oriented had low *Z*-values (**Fig. 6F**). To distinguish whether ectopic protrusions nevertheless had any preferred orientation, *Z*-values were also calculated for just the ectopic protrusions **(Fig. 6G)**. A high *Z*-value in these cases indicates that the ectopic protrusions are aligned, just not toward the oocyte. A subset of composite circular histograms illustrates some of the phenotypes identified by this analysis **(Fig. 6H)**. This analytical approach characterized protrusion phenotypes that were too subtle to detect by measuring migration defects alone. We conclude that GAPs restrict the frequency and range of ectopic protrusions in wild-type border cell clusters.

### Circularity and convex hull area provide global morphology measurements

Our stringent criteria for protrusion detection focused on large protrusions typically found in leading cells while excluding other extensions such as small blebs or clusters of filopodia. Complementary measurements of convex hull area and circularity (**Fig. 7A**) capture additional morphological features, including smaller protrusions and overall cluster shape, together providing a comprehensive morphological profile for each genotype. Convex hull area is the area of a polygon drawn from the most extended points. Circularity is a ratio of the area to the perimeter and measures how round an object is. Circularity ranges from 0 (not circular) to 1 (perfectly circular). We compared these measurements to the protrusion counts from the radial and rose plot analyses. As expected, convex hull area correlated positively with the number of protrusions (**Fig. 7B**), whereas circularity negatively correlated with protrusions (**Fig. 7C**). However, the correlations were imperfect, indicating that convex hull area, circularity, and protrusiveness capture distinct morphological features. We also compared circularity (**Fig. 7D**) and convex hull area (**Fig. 7E**) across the RNAi lines.

**Figure 7:**
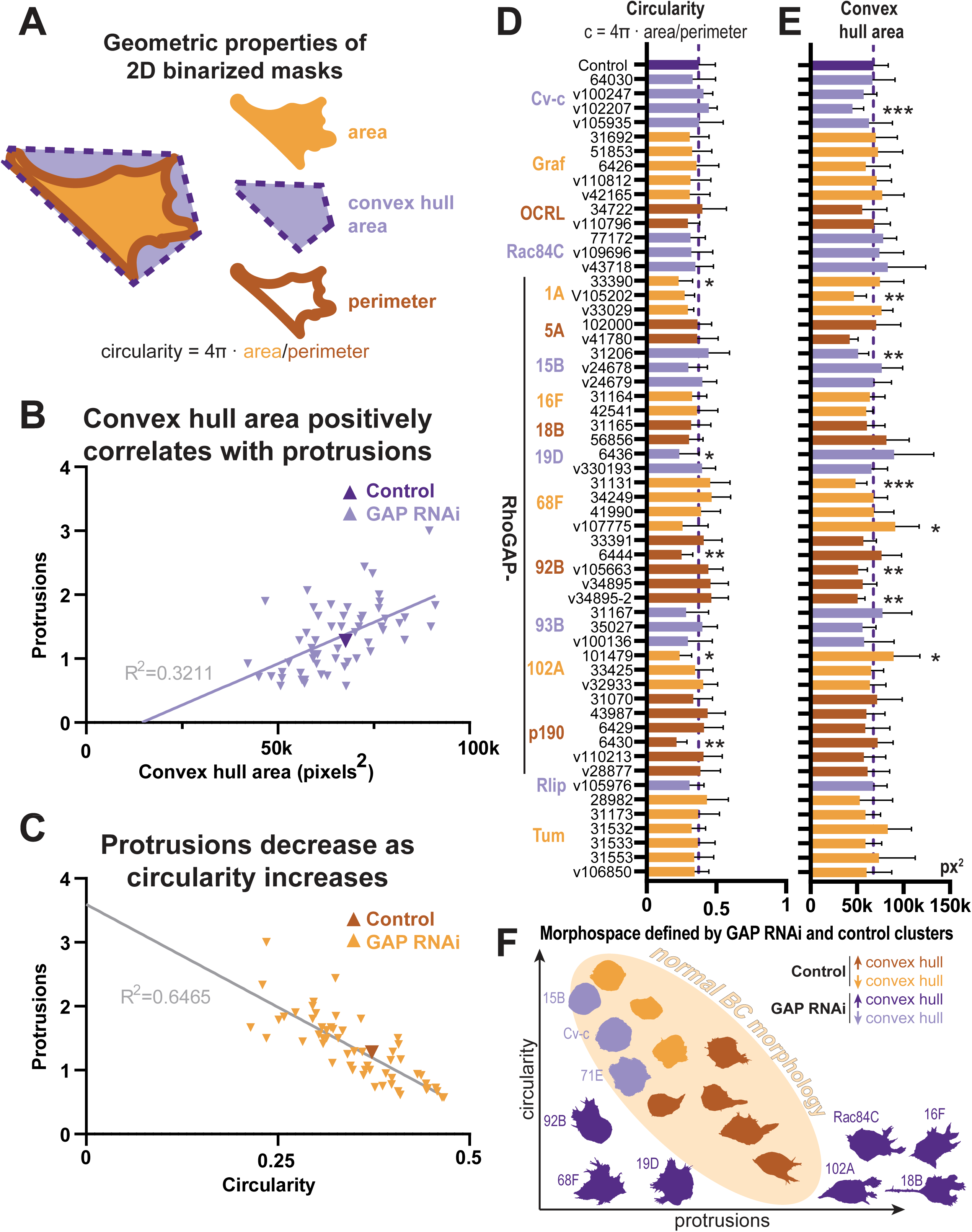
Circularity and convex hull area provide global morphology measurements. (A) Circularity and convex hull area quantify global geometric properties of 2D border cell images. Convex hull area captures large and small extensions. **(B)** Convex hull area correlates positively with the number of total protrusions. **(C)** Protrusions decrease as circularity increases, demonstrating the inverse relationships between these two properties. **(D)** Circularity is modulated by RhoGAPs. **(E)** Convex hull area is modulated by RhoGAPs. **(F)** Summary model of morphological measurements showing the range of wildtype circularity, protrusiveness, and convex hull area, with GAP knockdowns lying outside this wildtype morphological space. Bars show the mean ± SD. Significance reported using a Bayesian ANOVA. Bayes Factor BF_10_ >3 (*), BF_10_>10 (**), BF_10_ >30 (***).

To integrate all of the measured effects on cluster morphology, we graphed circularity on the y-axis and the number of protrusions on the x-axis and color-coded convex hull area. Wildtype clusters populated a limited range of phenotypes, from round to multiprotrusive, defining a characteristic morphological space **(Fig. 7F)**. By contrast, RhoGAP knockdown clusters dramatically extended this morphological space beyond the normal range, in multiple dimensions. Some knockdowns (RacGAP84C, RhoGAP102A, RhoGAP18B, RhoGAP16F) led to excessive protrusions and increased convex hull area, which pushed the boundaries of the protrusion axis beyond the wildtype range. Other GAP knockdowns introduced abnormal morphologies, including filopodia and blebs that increased the convex hull area through non-traditional protrusive structures. Jointly, these perturbations reveal that RhoGAP activities contribute to defining the normal border cell morphospace.

### Effects of RhoGAPp190 overexpression on protrusions and convex hull area

RhoGAPp190 is conserved from flies to mammals and negatively regulates Rho to control cytoskeletal organization. In cultured mammalian cells, p190 promotes adherens junction formation (Wildenberg et al., 2006), and a role in collective cell migration has been proposed based on p190 localization to cell-cell junctions (Zegers and Friedl, 2014). However, functional analysis of p190 in collective cell migration in vivo has not been reported.

We compared four conditions: control (LacZ), p190 RNAi knockdown, and p190 overexpression using either full-length protein or a truncated version (p190*) lacking the C-terminal RhoGAP catalytic domain but retaining all other identified domains, including the N-terminal GTPase-binding domain and middle regulatory domains. Low magnification images of stage 10 egg chambers revealed overall cluster positioning and morphology **(Fig. 8A-D)**, while high magnification highlighted detailed morphology **(Fig. 8E-H)**.

**Figure 8:**
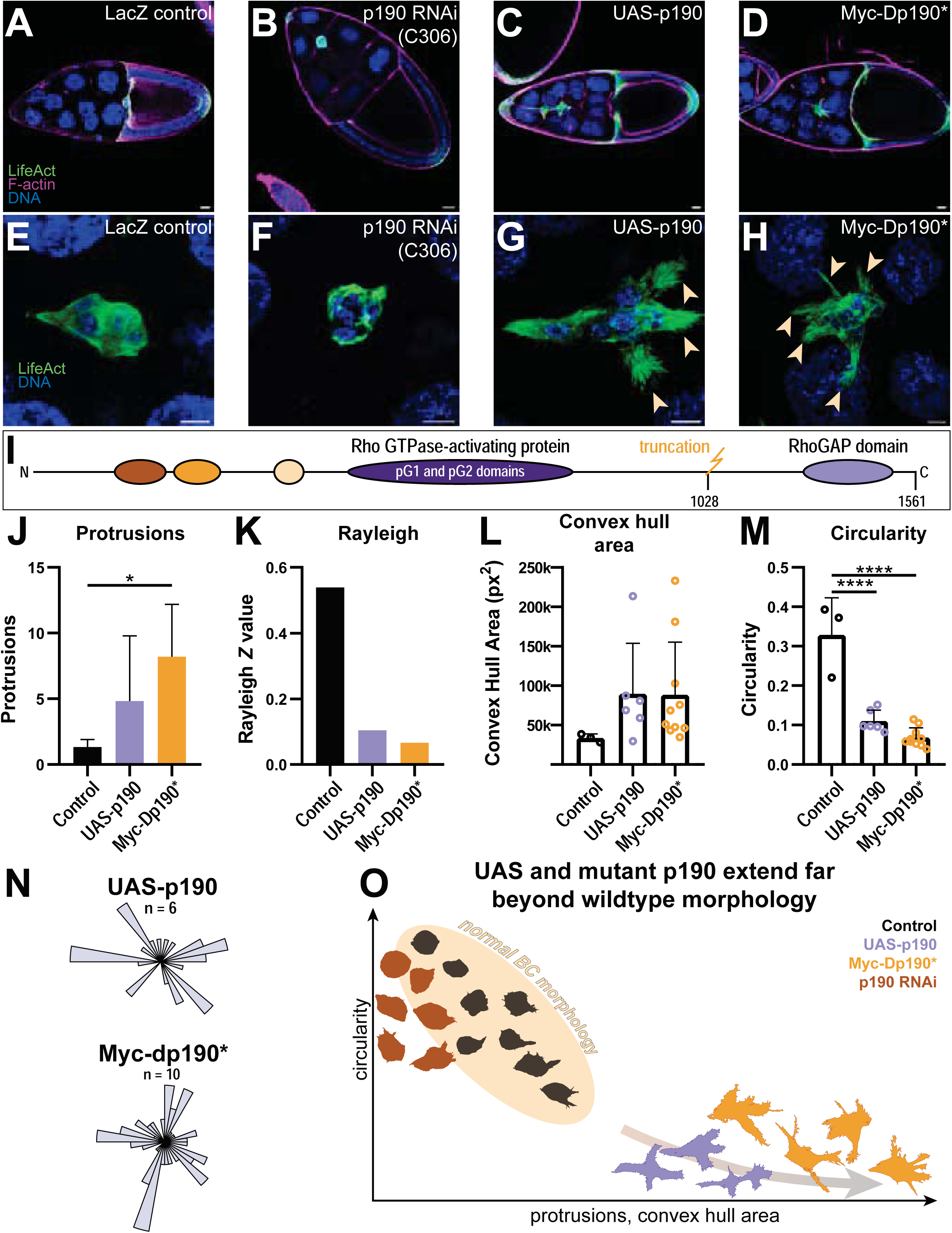
Effects of RhoGAPp190 overexpression on protrusions and convex hull area. (A) 20x image of control border cell clusters at stage 10. **(B)** c306Gal4>RhoGAPp190 RNAi at stage 10. **(C)** UAS-RhoGAPp190 (overexpression) at stage 10. **(D)** RhoGAPp190 truncated mutant (p190*) at stage 10. Scale bar = 20 𝜇m. **(E-H)** 63x representative images of the LacZ control **(E)**, p190 RNAi **(F)**, p190 overexpression **(G)**, and p190*. **(H)**. Multiple protrusions are labeled with yellow arrowheads. Scale bar = 10 𝜇m. **(I)** Protein domain schematic of RhoGAPp190 and p190*, showing the truncation at amino acid 1028 that removes the RhoGAP catalytic domain. **(J)** Protrusion counts of border cell clusters increased for UAS-p190 and - p190*. **(K)** Rayleigh *Z*-test values for protrusions directionality. **(L)** Convex hull area measurements, with each circle representing a single border cell cluster. **(M)** Circularity of border cell clusters. **(N)** Rose plots for the UAS-p190 and p190* border cell clusters. Replicates (n) are shown. **(O)** Morphospace plot showing the divergence of clusters expressing p190RNAi, UAS-p190, or p190* from the normal range of border cell morphologies. *p<0.05, **p<0.005, ***p<0.0005, ****p<0.0001. Mean ± standard deviation. One-way ordinary ANOVA with Dunnett correction for multiple comparisons.

Control clusters exhibited typical compact morphology (**Fig. 8A, E**, average 1.2 protrusions per cluster). p190 RNAi caused clusters to become rounder **(Fig. 8B, F)**, resembling clusters expressing constitutively active Rho **(Fig. 1G-I)**. In contrast, p190 overexpression produced dramatically splayed morphology with reduced cell-cell contacts within the cluster and large, long protrusions extending in all directions **(Fig. 8C, G)**. A strikingly similar phenotype was observed upon overexpression of a truncated protein (hereafter p190*), which lacks the GAP domain but retains all other protein-protein interaction domains **(Fig. 8D, H and I)**. Quantitative morphological analysis revealed significant differences across conditions **(Fig. 8J-O)**. Overexpression of full-length p190 increased protrusions from 1.2 to 5, and p190* overexpression increased protrusions further to 8 **(Fig. 8J)**. Protrusion directionality, measured by the Rayleigh *Z*-test, decreased from ∼0.5 in control to ∼0.1 with p190 overexpression and ∼0.05 with p190* overexpression, indicating increasingly random protrusion orientation **(Fig. 8K)**. Convex hull area, reflecting overall cluster spread, increased approximately 2.5-fold in both p190 and p190* overexpressing clusters compared to control **(Fig. 8L)**. Circularity decreased from ∼0.3 in control to ∼0.1 with p190 overexpression and ∼0.07 with p190* overexpression, reflecting the transition from compact to highly irregular morphology **(Fig. 8M)**. Rose plots show the angular distribution of protrusions in p190 and p190* overexpressing clusters, confirming the loss of directional bias **(Fig. 8N)**.

Plotting these measurements in morphospace—with protrusion number on the x axis and circularity on the y—revealed that p190 overexpression pushed clusters far outside the normal morphospace occupied by wild-type cells and in the opposite direction compared to p190 RNAi knockdown **(Figure 8O)**. The high protrusion numbers in clusters overexpressing p190* drove them even further into abnormal morphospace (8N, see discussion). These gain-of-function phenotypes were remarkably similar to those described for knockdown of myosin II in border cell clusters (Mishra et al., 2019).

### Effects of RhoGAPp190 knockdown on ɑ-catenin and myosin abundance and localization

To explore the mechanistic basis of p190 function in border cells, we examined the localization and abundance of α-catenin, a key component of adherens junctions that links cadherins to the actin cytoskeleton, and myosin II. We compared control and p190 RNAi-expressing clusters at three stages: during detachment from the anterior follicle epithelium, mid-migration between nurse cells, and post-migration after reaching the oocyte. We visualized non-muscle myosin II using Sqh-mCherry, a fusion of the non-muscle myosin II regulatory light chain Spaghetti squash (Sqh), expressed under the *sqh* regulatory sequences **(Fig. 9A-F)** and α-catenin using anti-DCAT-1 antibody **(Fig. 9A’-F’)**.

**Figure 9:**
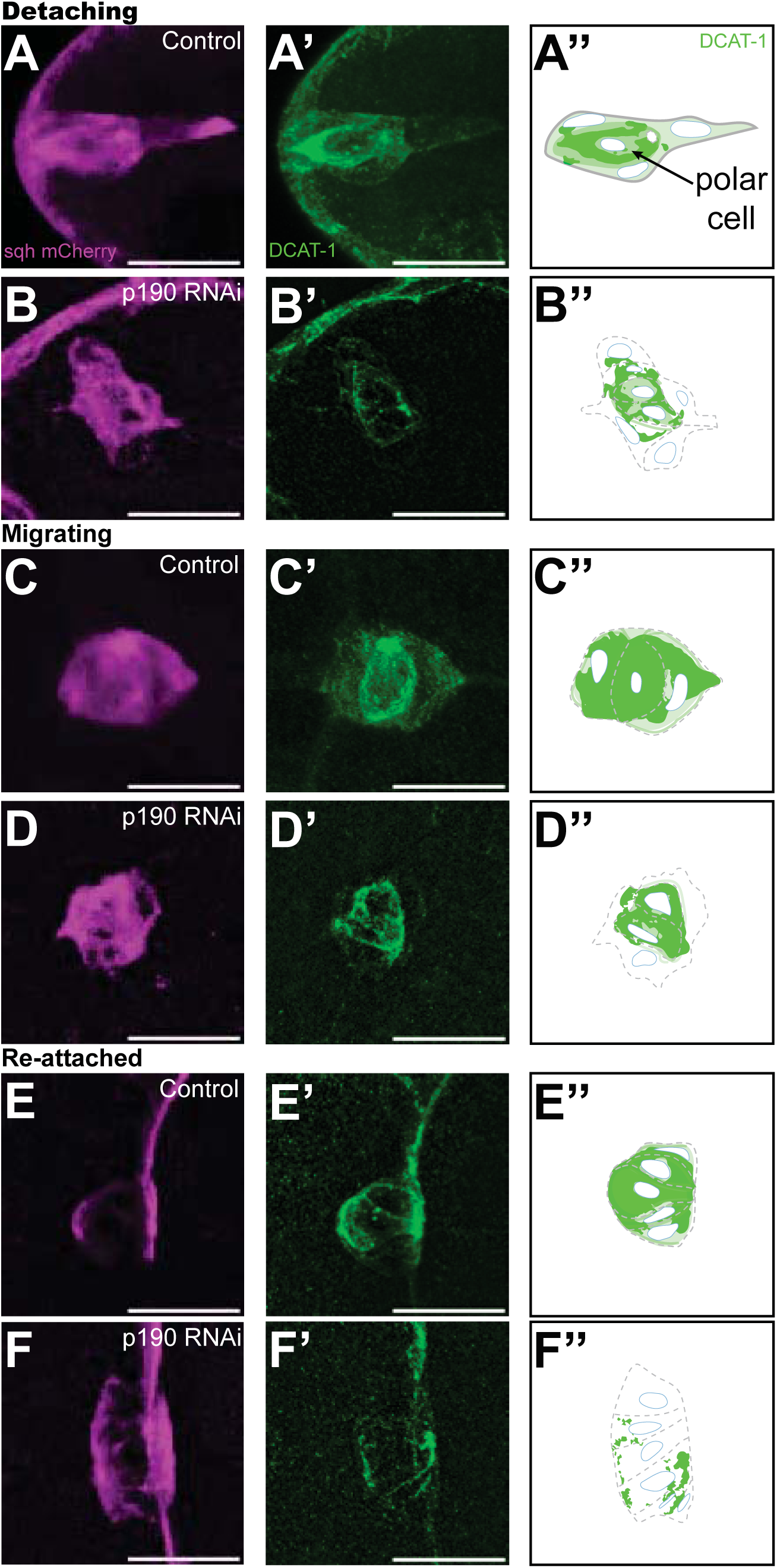
Effects of RhoGAPp190 knockdown on ɑ-catenin and myosin abundance and localization. (A-A”) Control cluster during detachment showing the distribution of Sqh-mCherry (non-muscle myosin II light chain) **(A)**, anti-alpha-catenin antibody staining (DCAT-1) **(A’)**, and a schematic showing DCAT-1 localization **(A’’)**. **(B-B”)** The same markers in a p190RhoGAP RNAi-expressing cluster during detachment. **(C-C”)** A control cluster mid-migration. **(D-D”)** A p190RhoGAP RNAi-expressing cluster mid-migration. **(E-E”)** A post-migratory control cluster. **(F-F”)** A p190RhoGAP RNAi-expressing cluster post-migration.

In control clusters, α-catenin localized to cell-cell junctions throughout migration **(Fig. 9A’, C’, E’)**. p190 RNAi caused a notable reduction in α-catenin protein levels across all stages examined **(Fig. 9B’, D’, F’)**. Myosin II (sqh-mCherry) showed similar distribution in control and p190 RNAi clusters during detachment and post-migration **(Fig. 9A-B, E-F)**. However, at mid-migration, p190 RNAi clusters exhibited elevated myosin II levels compared to control **(Fig. 9D vs. C)**, suggesting increased actomyosin contractility during active migration when p190 is depleted, consistent with the morphological phenotypes including rounded and split clusters, increased circularity, decreased protrusions.

## Discussion

Our detailed analyses of RhoGAPp190 illustrate how systematic phenotyping can guide mechanistic investigation. p190 knockdown phenocopied Rho^V14^ expression, while p190 overexpression resembled myosin II loss, supporting a model in which p190 restrains Rho-ROCK-myosin signaling to maintain the proper balance of contractility and cell-cell adhesion. The observation that p190* overexpression increases protrusiveness, like p190 overexpression, rather than producing an opposite effect or no effect, is surprising given that p190* lacks GAP catalytic activity. This suggests p190’s function extends beyond simple GTP hydrolysis. One interpretation is that p190 serves as a spatial organizer of Rho signaling complexes, and both overexpression conditions disrupt the normal stoichiometry or localization of these complexes. Alternatively, p190* may sequester a negative regulator, leading to hyperactivation of the endogenous full-length protein. The distinct effects of p190 loss-of-function (rounder, less protrusive) versus p190 gain-of-function (splayed, hyperprotrusive) demonstrate that precise regulation of p190 activity levels and/or localization is critical for normal cluster morphology and cell-cell adhesion. This example demonstrates how the comprehensive phenotypic profiling developed for the screen will generally enable deeper and more sensitive characterization of genes and proteins that control border cell morphology.

Zegers and Friedl (Zegers and Friedl, 2014) proposed that RhoGAPp190 would promote adhesion between collectively migrating cells, and the current study provides the first demonstration of that function *in vivo*. The reduced α-catenin levels in p190-depleted clusters, combined with their rounder morphology and reduced protrusiveness, indicate that p190 maintains proper junctional organization. The elevated myosin II in mid-migration p190 RNAi clusters is consistent with p190 normally restraining Rho-ROCK-myosin signaling. Together, these findings support a model in which p190 negatively regulates Rho-ROCK-myosin signaling to balance contractility, protrusion dynamics, and cell-cell adhesion within the migrating cluster.

Our systematic analysis reveals that beyond p190, most or all Drosophila RhoGAPs are expressed in border cells and functionally required for normal morphology or migration. Importantly, no single expression profiling technique detected all RhoGAPs; rather, the pattern emerged by integrating datasets with different sensitivities. More sensitive methods like bulk RNAseq of purified border cells detected lowly expressed genes missed by microarray, while highly expressed genes were detected across multiple platforms. This highlights the value of complementary approaches for comprehensive gene expression analysis in rare cell populations.

Expression level did not predict phenotypic severity. RhoGAP19D showed the highest expression but no detectable migration defect, possibly reflecting insufficient RNAi-mediated knockdown of abundant transcripts. Conversely, RacGAP84C and RhoGAP92B, detected at low levels, produced clear migration defects with multiple independent RNAi lines. This suggests either that RNAi is more effective against lowly expressed genes or that cells are particularly sensitive to loss of certain GAPs even when modestly expressed.

Migration assays proved surprisingly insensitive for detecting phenotypic effects. A striking example is constitutively active Rho, which causes dramatic morphological changes yet only mild migration defects, likely reflecting the plasticity of migration modes. To capture phenotypic complexity more sensitively, we developed automated image analysis tools that objectively quantify circularity, convex hull area, protrusion number, and directionality. This revealed that different RhoGAP knockdowns produce distinct morphological signatures, suggesting specialized rather than simply redundant functions.

A key insight emerged from plotting morphological parameters: wild-type border cells occupy a defined “morphospace,” and many RhoGAP knockdowns pushed clusters outside this normal range. This reveals that GAPs actively constrain cell morphology within functional boundaries. The normal morphospace boundaries likely reflect both external constraints from surrounding tissue as well as internal factors. External constraints from confinement within the egg chamber would apply equally to wild-type and RNAi-treated clusters, since the RNAi was specific to the border cells, which may limit the morphospace that the knockdown clusters can occupy. Evidence for external constraints includes the dramatic lamellipodia border cells produced when E-cadherin is knocked down specifically in nurse cells, opening larger spaces for broader protrusions than normally occur (Dai et al., 2020). Nevertheless we observed dramatic deviations from normal morphology, especially in p190 and p190* overexpression.

The complexity we observe—22 GAPs regulating 5 GTPases in 6-8 cells—suggests RhoGTPase activities are sculpted by overlapping GAP activities into precise spatiotemporal patterns. Our systematic approach in border cells provides a roadmap for dissecting this complexity in other migratory cell types and understanding how cells orchestrate cytoskeletal regulation during collective migration.

## Materials and methods

### Software

- <u>Ubuntu</u> (version 24.04.2)
- The Broad Institute <u>Drop Seq Tools</u> (version 2.4.0)
- R (version 4.5.1)
- <u>RStudio</u> (version 2025.05.1+513) with the following packages: <u>Seurat</u> (5.3.1), SeuratObject (5.2.0), ggplot2 (4.0.0), tidyverse (2.0.0), and patchwork (1.3.2).
- Python 3.13.15 and Spyder 6.0.7 using the following packages: os, numpy (2.1.3), pandas (2.2.3), skimage (0.25.0), scipy (1.15.3), matplotlib (3.10.0), pathlib, aicspylibczi (3.3.1), PIL (11.1.0), and tkinter.
- <u>ImageJ2</u> by (Rueden et al., 2017)
- <u>JASP stats</u> (version 0.95.4.0) by JASP team (JASP Team, 2025)
- <u>Prism</u> (version 8.0.1) by GraphPad
- <u>Adobe Illustrator</u> 2026 by Adobe
- <u>Zen</u> by Zeiss

### Methods

#### *Drosophila* egg chamber single-cell RNA sequencing

*upd-Gal4, UAS-DsRed.nls; slbo>Lifeact-GFP* flies were used for single-cell RNA sequencing to label border cells (GFP) and polar cells (DsRed). Ovaries were dissected in Schneider’s media with 0.1 mg/mL of insulin and kept on ice. Late stage 9/early stage 10 egg chambers were teased away from other stages with a dissecting needle in Schneider’s media on a 4% agarose pad, taking care to avoid late stage 10 and older. Dissociation was performed in 100 μL of elastase with gentle trituration for 5 minutes. Dissociation was quenched with 300 μL of 80% Schneider’s medium to 20% FBS. Cells were then rinsed of elastase with .01% BSA in PBS 3 times. The resulting suspension was filtered through a 40–70 μm cell strainer, and all steps were performed in the cold to preserve cell viability and RNA integrity. Single-cell transcriptomes were generated using Drop-seq technology (Macosko et al., 2015) (**Figure S1**). Libraries were prepared according to standard Drop-seq protocols and sequenced on an Illumina HiSeq 2500 platform. Three biological replicates of 500 egg chambers each for a total of ∼1500 egg chambers were analyzed.

#### RNA sequencing analysis in Shell

Data were processed according to the recommended protocol by the Broad Institute, which provides a publicly available set of DropSeq tools on <u>GitHub</u>. These were installed using `setup_dropseq.sh`. Drop-seq-tools version 2.4.0 was used for this analysis.

##### Download reference genome

The *Drosophila melanogaster* reference genome (dmel_r6.63_FB2025_02) was downloaded from FlyBase using `download_reference.sh` and saved to a results directory alongside the references directory. The reference genome was enhanced with the experimental transgenes (dsRedExpress, LifeActGFP) using `enhance_references_with_transgenes.sh`.

##### Combine sequencing lanes for DropSeq sample

Using the experimental data collected from stage 10 *Drosophila* ovaries, the individual lanes from each replicate (d1, F1, F2) were merged using the `combine_lanes.sh` script. For each sample, it takes the multiple .fastq.gz files and combines them into two .fastq.gz files, one for cell barcodes and one for transcripts. These larger files were then used for downstream analysis.

##### DropSeq pre-processing

FASTQ files were converted into unmapped binary alignment map (BAM) files. Cell and molecular barcodes were tagged, and the BAM file was filtered. SMART adapters and polyA regions were trimmed, and then the file was converted back into a FASTQ file.

##### STAR alignment

Pre-processed FASTQ files for each sample were aligned to the reference genome (dmel_r6.63_FB2025_02) using Spliced Transcripts Alignment to a Reference (<u>STAR,</u> <u>version 2.7.11b</u>. Alignments were performed with the following parameters: output format was set to coordinate-sorted BAM files with unmapped reads included. The number of report alignments (NH), hit index (HI), alignment score (AS), number of mismatches (nM), edit distance (NM), and string encoding mismatched and deleted reference bases (MD) were retained for downstream analysis. Filtering was performed using splice junction-based criteria, allowing up to 20 multi-mapping locations per read. Mismatch filtering permitted up to 999 total mismatches per read, but required that no more than 4% of mismatches constituted the total read length. Splice junction parameters were set based on the *Drosophila* gene structure, with a minimum intron length of 20 bp, maximum intron length of 1,000,000 bp, maximum gap between paired-end mates of 1,000,000 bp, minimum overhang of 8 bp for unannotated and 1 bp for annotated junctions, and a splice junction score of 1. These were optimized for single-cell RNA sequencing data and *Drosophila*-specific genomic figures. These steps were performed using `run_star_alignment_with_transgenes.sh`.

##### Create Digital Gene Expression Matrix

The STAR-aligned data was sorted by query name and then merged to recover cell and molecular barcodes. Gene exon tags were also added during this step, and the final gene expression data was organized into a Digital Gene Expression (DGE) matrix. These steps were jointly performed using `annotate_aligned_files_with_transgenes.sh`. This DGE matrix was then used in subsequent analysis using Seurat.

#### Seurat RNA sequence analysis in RStudio

The DGE matrix was further processed in RStudio using Seurat. DGE files (.txt.gz) for each sample (d1, F1, F2) were read into R and converted to individual Seurat objects with minimum thresholds of 3 cells per feature and 200 features per cell. The three samples were then merged with sample-specific cell IDs to enable batch tracking. Quality control metrics were calculated for each cell, including the number of detected genes (nFeature_RNA), total unique molecular identifier (UMI) counts (nCount_RNA), and percentage of mitochondrial reads. QC distributions were visualized using violin plots (Fig. S2A - S2C). Cells were filtered to keep only high-quality cells meeting the following criteria: 500-4000 genes per cell, less than 10% mitochondrial reads, and 1,000-15,000 UMIs per cell. The filtered dataset was log-normalized using the standard Seurat method (log[count/total counts per cell × 10,000 + 1]). Dimensionality reduction was performed using Principal Component Analysis (PCA), and the top 20 principal components were selected. Cell clustering was performed using the Leiden algorithm (algorithm = 4) with k.param = 8 for nearest neighbor identification and a resolution of 2.0. UMAP dimensionality reduction was performed using the top 20 PCs with a minimum distance parameter of 0.1. Cluster-specific marker genes were identified using the FindAllMarkers function with only positive markers retained. The processed Seurat object containing the filtered expression matrix, cluster assignments, and UMAP coordinates was saved for downstream visualization and analysis.

#### Plotting RNAseq data in RStudio

Using the final Seurat object, border cells were selected by filtering for Slbo-, LifeActGFP-, and Fruitless (Fru)-positive cells. Polar cells were selected by filtering for dsRedExpress. Rho-GTPase and -GAP genes were quantified for the border and polar cell populations. These expression patterns were compared side-by-side and sorted by border cell expression. Violin plots are additionally provided in Figure S2. The border and polar cell populations were also plotted onto the previously-created UMAP, showing that clustering was insufficient to identify these cell types. These steps were all performed using the script `plot_Seurat.R`.

#### RhoGAP RNAi screen

UAS RNAi fly stocks for all known RhoGAPs were ordered from the Bloomington Drosophila Stock Center (BDSC) and the Vienna Drosophila Resource Center (VDRC) unless they. See table S1 for RNAi lines used. Five to seven virgin females (genotype: c306-Gal4/c306-Gal4; slbo>LifeActGFP/Cyo; +/+) virgin females were crossed with three to four males carrying UAS-RNAi constructs. Crosses were grown at 18°C, and adults were flipped to new food vials every two days. Eight to ten female progeny and two to three male progeny were fattened at 30°C in heavily yeasted vials for 72 hours. Ovarioles were dissected in Schneiders Media (ThermoFisher, catalog #21720001) supplemented with 20% fetal bovine serum and fixed for 15 minutes in 4% paraformaldehyde (Electron Microscopy Sciences) diluted in phosphate-buffered saline (PBS).

Ovarioles were washed three times for 15 minutes in PBS+0.2% tritonX-100 (PBST) before 30 min incubation in Hoechst (1:1000 dilution of a 10mg/ml stock), and 1:400 dilution of Phalloidin Atto647N (Sigma-Aldrich, catalog# 65906-10NMOL). Egg chambers were washed two times for 15 min in PBS before mounting in VectaShield (Vector Laboratories, catalog #H-1000) on a #1.5 coverglass and imaged with a Zeiss LSM800 confocal microscope using the PlanAPO 20x 1.2NA and PlanAPO 40x 1.4NA objectives. All stage 10 egg chambers from each dissection were manually counted for migration completion by stage 10 using the 20x. Additionally, 20 border cells were randomly selected for morphology analysis after imaging at 40x.

#### Circularity and convex hull area analysis of GAP knockdown border cell clusters

Border cell clusters were pre-processed in FIJI/ImageJ to select for the membrane channel, maximum project the cluster, and convert the image to grayscale. A macro was used to perform this on all images using the script `Unmerge_MaxProj_SaveTIF.ijm`. These preprocessed 2D images were then binarized and segmented in Python using scikit-image. The clusters were segmented using Li thresholding, which provided the best segmentation of filopodia and small-scale details. This segmented object was filled, small objects were removed, and the largest object was selected, which created a 2D image of the border cells. Clusters that were detaching or reattaching were excluded. RNAi lines with fewer than 7 clusters were excluded from morphological analysis. These segmented images were then measured for perimeter, area, and circularity using numpy, and convex hull area using scipy.spatial. All images were iterated through and measured using this same method, and the results were saved to csv files for each genotype. These analyses were all performed using `global_morphology.py`. The csv files were then compiled into a single file using `combine_global_morphology_files_csv.py`.

#### Border cell re-orientation using Python GUI

Each cluster required reorientation in the direction of migration to determine ectopic versus lead protrusions. To optimize this process, a custom graphical user interface (GUI) was written in Python to open the 40x and corresponding 20x image for each border cell cluster. These were displayed side by side. To set the angle, the user clicks once on the center of the border cell cluster in each 20x image and once in the direction of the oocyte. The angle of this vector was saved and used later in the radial and rose plot analyses to use as the rotation for each cluster. This GUI is available in `re-orientation_GUI.py`.

#### Radial and rose plot analyses to quantify protrusions and directionality

To quantify protrusion number and angular distribution, the binarized and segmented border cell cluster images were analyzed using custom Python scripts. For each cluster, the boundary contour was extracted and converted to polar coordinates centered on the cluster centroid. The radial distance from the centroid to the boundary was measured at 360 angular positions (1° bins), creating a radial profile. Each cluster was rotated using the orientation angle determined in the GUI, positioning the direction of migration towards the oocyte at 0°. Protrusions were identified as peaks in the radial profile using two thresholds implemented through scipy.signal.find_peaks. First, a height threshold was set at 120% of the mean cluster radius to identify radial extensions that extend substantially beyond the average cell boundary. Second, a prominence threshold was set at 50% of the mean radius. Prominence measures how much a peak stands out relative to its immediate surroundings. Quantitatively, it is the vertical distance between the peak and the highest contour line that could encircle the peak without encircling a higher peak. This criterion filtered out minor boundary irregularities or plateaus, while retaining genuine cellular extensions. Together, these thresholds provide a robust, quantitative definition of a protrusion.

Lead and ectopic protrusions were classified by radial position; ±20° from the 0° vector (oocyte) was selected for lead protrusions, while all other directions were considered ectopic. For each genotype, a composite rose plot was created by aggregating protrusions by angle into 36 angular 10° bins. Rose plot values were normalized by dividing the number of protrusions by the number of replicates in each genotype. This normalization allowed for direct comparison across genotypes with different sample sizes. Directional bias in protrusion orientation was quantified using the Rayleigh test for circular uniformity (*Z*-test). The Rayleigh test calculates a resultant vector (*Z*) from the angular distribution of the protrusions in the rose plot, where *Z* ranges from 0 (uniform distribution, no directional preference) to 1 (all protrusions are oriented in the same direction). Z was calculated for pre-orientation (Figure S4D), oriented, and only ectopic protrusion angles. All of these analyses were performed in a custom Python script, `radial_analysis_rose_plots.py`.

#### Fly stocks, immunofluorescence, and imaging for RhoGTPase and RhoGAPp190 experiments

##### RhoGTPase experiments

The *Gal4* driver *slboGal4/cyo; slbo>LifeActGFP* was crossed with UAS-transgenes to express LacZ (control) or constitutively active GTPases in the border cells. These lines are reported in Table S1. Constitutively active GTPases included UAS-Cdc4^V12^, UAS-Rac1^V12^, and UAS-Rho1^V14^. Progeny from these crosses were fattened at 30°C in yeast-supplemented vials for 24 hours before dissection of egg chambers. Samples were mounted in Vectashield (Vector Laboratories) before imaging with a confocal microscope.

##### UAS-RhoGAPp190 and UAS-RhoGAPp190* experiments

The *Gal4* driver *slboGal4/cyo; slbo>LifeActGFP* was crossed with UAS-LacZ, UAS-myc-RhoGAPp190, and UAS-myc-RhoGAPp190* (stocks listed in Table S1) to perform gain-of-function experiments. Progeny from these crosses were fattened at 30°C in yeast-supplemented vials for 24 hours before dissection of egg chambers. Samples were mounted in Vectashield (Vector Laboratories) before imaging with a confocal microscope.

##### RhoGAPp190 and α-catenin experiments

The *GAL4* drivers *c306-Gal4;+; sqh-mcherry* was crossed with UAS-LacZ or UAS-RhoGAPp190 RNAi (VDRC #110213) to express fluorescent myosin in control or p190 knockdown border cells. Progeny from these crosses were fattened at 30°C in yeast-supplemented vials for 72 hours before dissection of egg chambers. Immunostaining was performed following the protocol previously described in Mishra et al., 2019. The following primary antibodies were used: rat anti–α-catenin (DSHB, DCAT-1) at a dilution of 1:20, and rabbit anti-GFP (Thermo Fisher Scientific, G10362) at a dilution of 1:1000. Secondary antibodies conjugated to Alexa Fluor 488 or Alexa Fluor 568 (Thermo Fisher Scientific) were used at 1:400 dilution, and F-actin was stained with Phalloidin–Atto 647N (Sigma-Aldrich). Samples were mounted in Vectashield (Vector Laboratories) before imaging.

#### Statistical analysis and data presentation

##### Morphology statistics

Bayesian ANOVA analyses were carried out using JASP. Post hoc tests were performed with a null control correction. Bar graphs were plotted in GraphPad Prism showing the mean and standard deviation. Significance was reported using the following thresholds: Bayes Factor BF_10_ >3 (*), BF_10_>10 (**), BF_10_ >30 (***). Bayesian statistics were selected to quantify the probability that each genotype differed from the control, rather than doing 70+ hypothesis tests across each RNAi line. This avoided multiple testing penalties while still being rigorous about false positives.

##### Linear correlation plots

Linear regressions were performed in GraphPad Prism, with R^2^ values reported on each plot.

**Figure 8**: Ordinary One-way ANOVA analyses and Dunnett’s correction for multiple comparisons were carried out using GraphPad Prism. Graphical representations were produced in GraphPad Prism. Figures show the mean with standard deviation. Significance was reported using the following thresholds: p<0.05 (*), p<0.005 (**), p<0.0005 (***), p<0.0001 (****).

##### Data presentation

RNAseq UMAPs were created in RStudio. Binary masks and rose plot figures were created in Python using Matplotlib. Bar plots and correlation plots were created in GraphPad Prism. Figures were created in Adobe Illustrator 2026. Scripts for all analysis steps are available on <u>GitHub</u>.

## Data availability statement

Data are shown in the figures and supplemental files, and additional data files are available upon request. Scripts are provided on <u>GitHub</u>.

## Acknowledgments

We gratefully acknowledge the Bloomington Drosophila Stock Center and the Vienna Drosophila Resource Center for Drosophila stocks, and Alba Torres Espinosa for Rac migration defect data. We thank Joshua Garcia, Victor Berkland, Amanda Cecil, Aaron Fulton and Nora Rudd for technical assistance. We acknowledge ClaudeAI for debugging and cleaning up scripts. We also thank Sreesankar Easwaran, Guangxia Miao, and Xiaoran Guo for help with egg chamber dissection and sample preparation for single cell RNAseq.

This work was supported by NIH grant R01GM073164 to D.J. Montell. E.G. Gemmill was supported by the National Institutes of Health T32 grant 1T32GM141846 to B.L. Pruitt and National Science Foundation predoctoral fellowship 2139319 to E.G. Gemmill.

## Author contributions

Designed experiments (A.K.M., J.A. M., J.P.C., E.G.G., V.R., D.J.M, K.S.K.), Performed experiments (A.K.M., J.A.M., J.P.C., E.G.G., V.L.), Designed and performed computational analyses (A.K.M., J.M., E.G.G.), Figures (E.G.G., J.P.C., A.K.M.) writing (E.G.G., V.L., K.S.K. D.J.M).

