## Supplemental Figures for "Systematic analysis of RhoGAP expression and function in border cell morphology and migration"

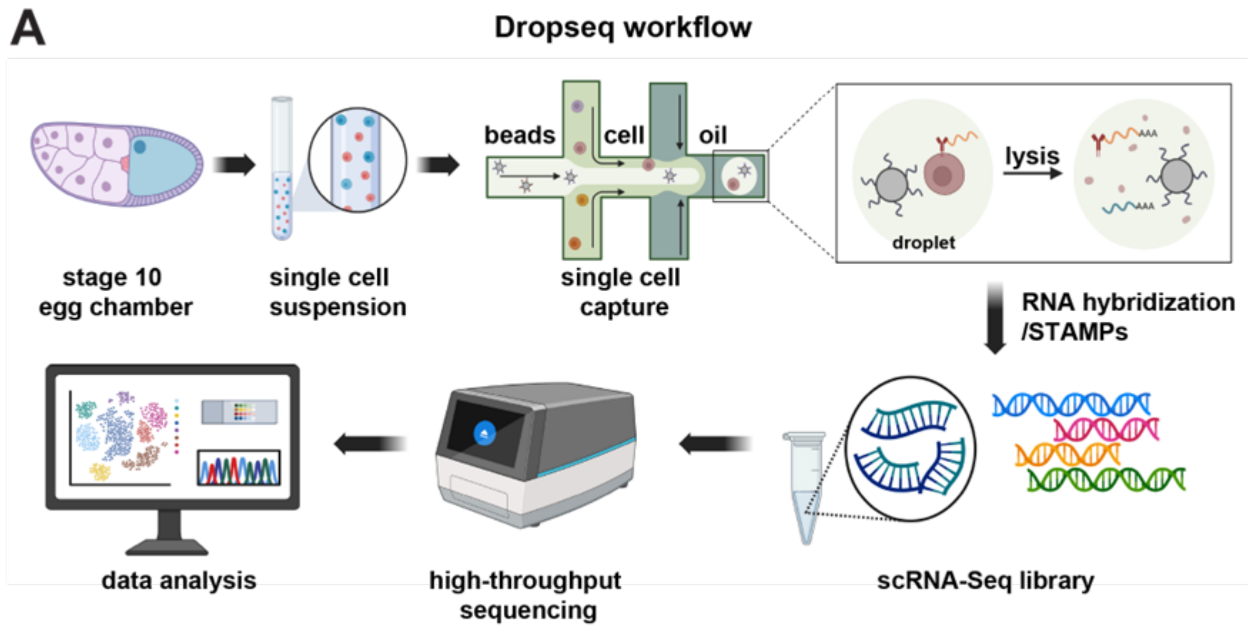


**Supplemental Figure 1: DropSeq workflow on stage 10 egg chambers. (A)** Stage 10 egg chambers are dissected, dissociated, and suspended, then processed according to standard DropSeq protocols as detailed in Methods. Created in BioRender. Mishra, A. (2026) https://BioRender.com/v79kui3

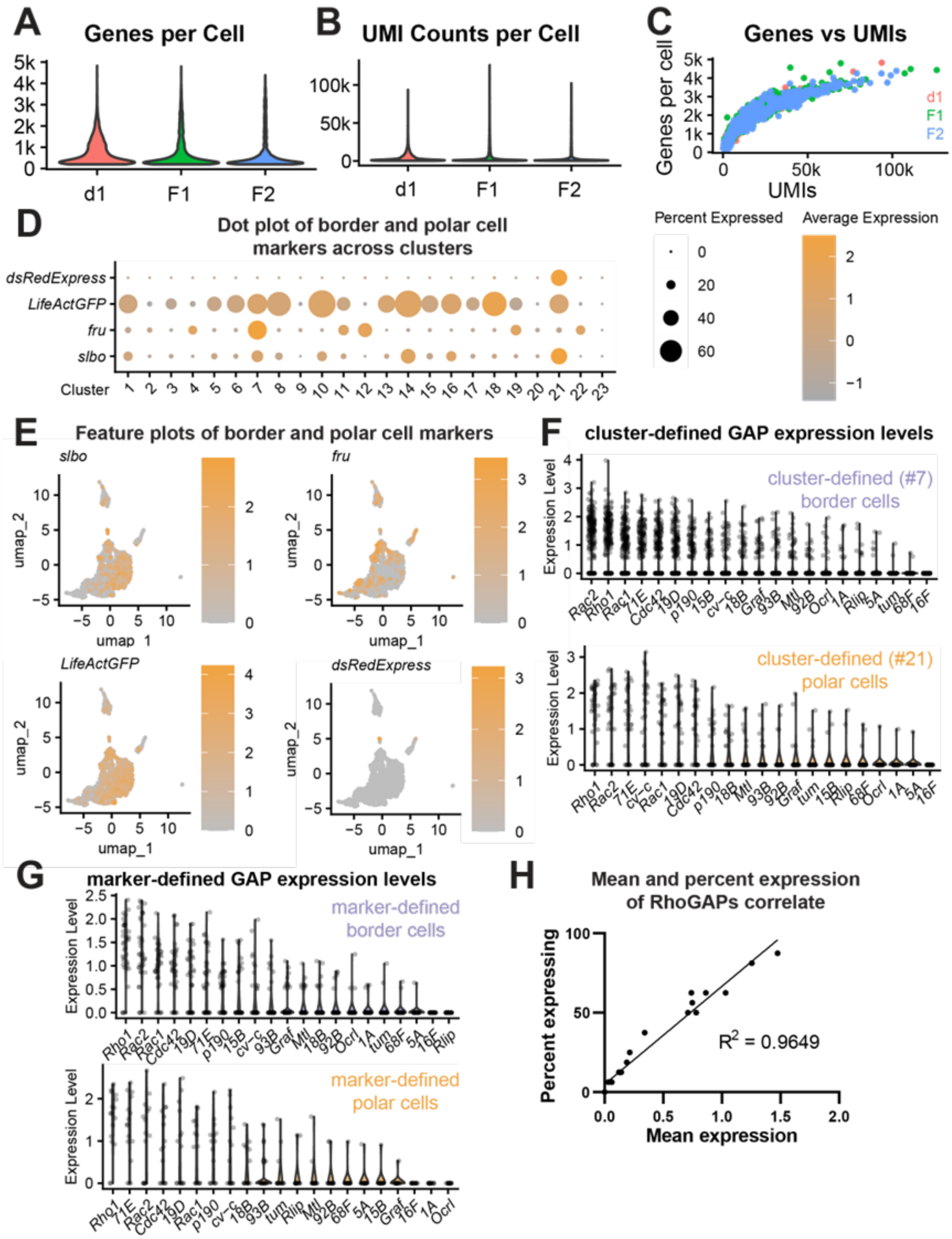


**Supplemental Figure 2: Single-cell RNAseq quality controls and clustering for the *Drosophila* ovary. (A)** The number of genes per cell before filtering cells in Seurat. **(B)** The number of unique molecular identifiers (UMIs) per cell before filtering **(C)** Elbow plot of the number of genes versus the number of UMIs. **(D)** Feature plot of dsRedExpress, marking polar cell populations. **(E)** Feature plots of slbo, fruitless (fru), and LifeActGFP. Cells expressing all three were selected to identify border cells. **(F)** Dot plot of the border and polar cell markers across the UMAP clusters. **(G)** GAP expression levels in border cells (top) and polar cells (bottom). Circles represent individual cells, with 16 polar cells and 36 border cells being quantified. **(H)** Mean and percent expression of RhoGAPs correlate significantly ($R^2$ = 0.9649).

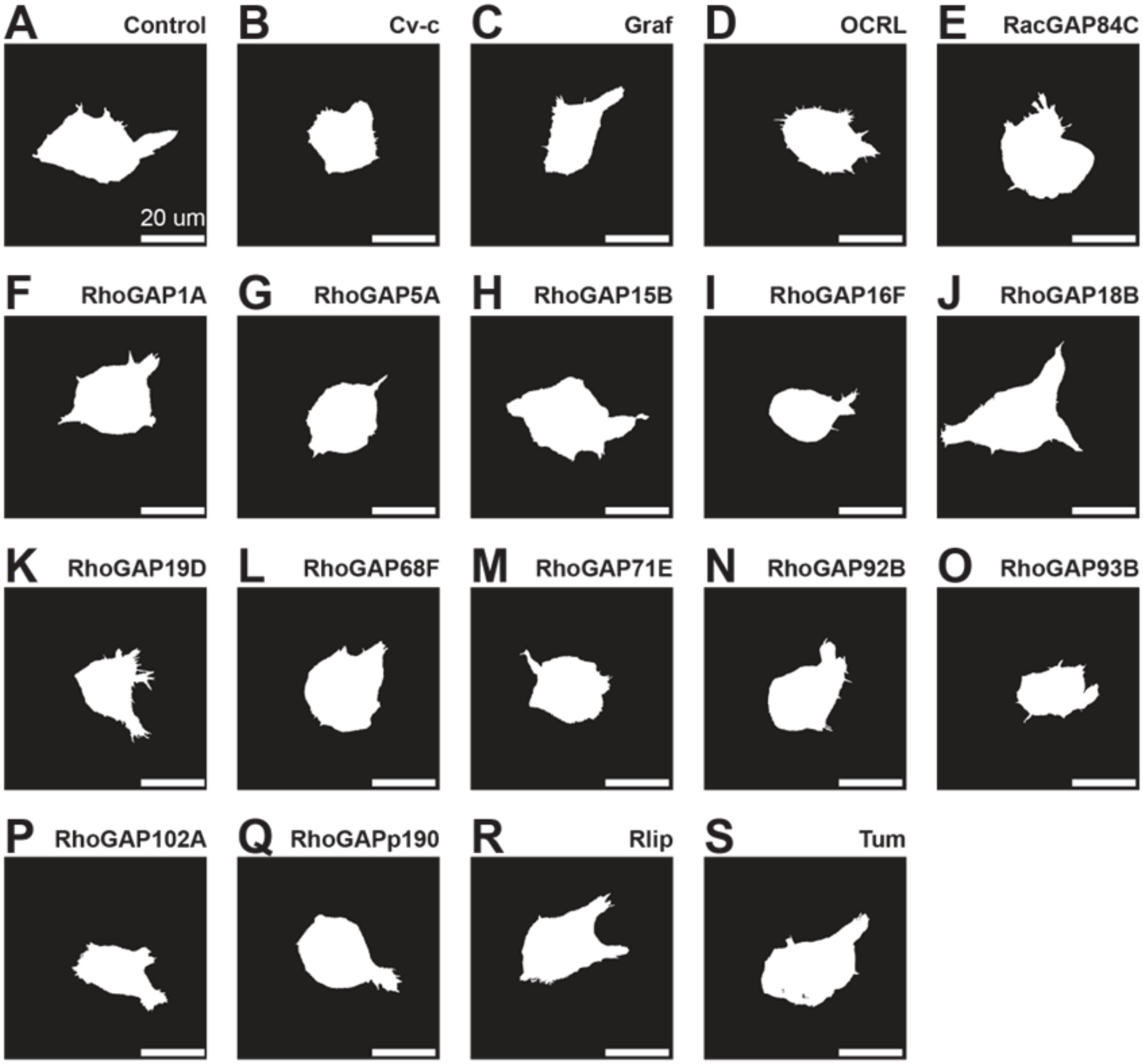


**Supplemental Figure 3: Representative binary masks of border cells with RhoGAP RNAi knockdowns for morphology analysis. (A)** Representative control cluster. **(B-S)** GAP RNAi knockdown clusters. Scale bar = 20 $\mu$m.

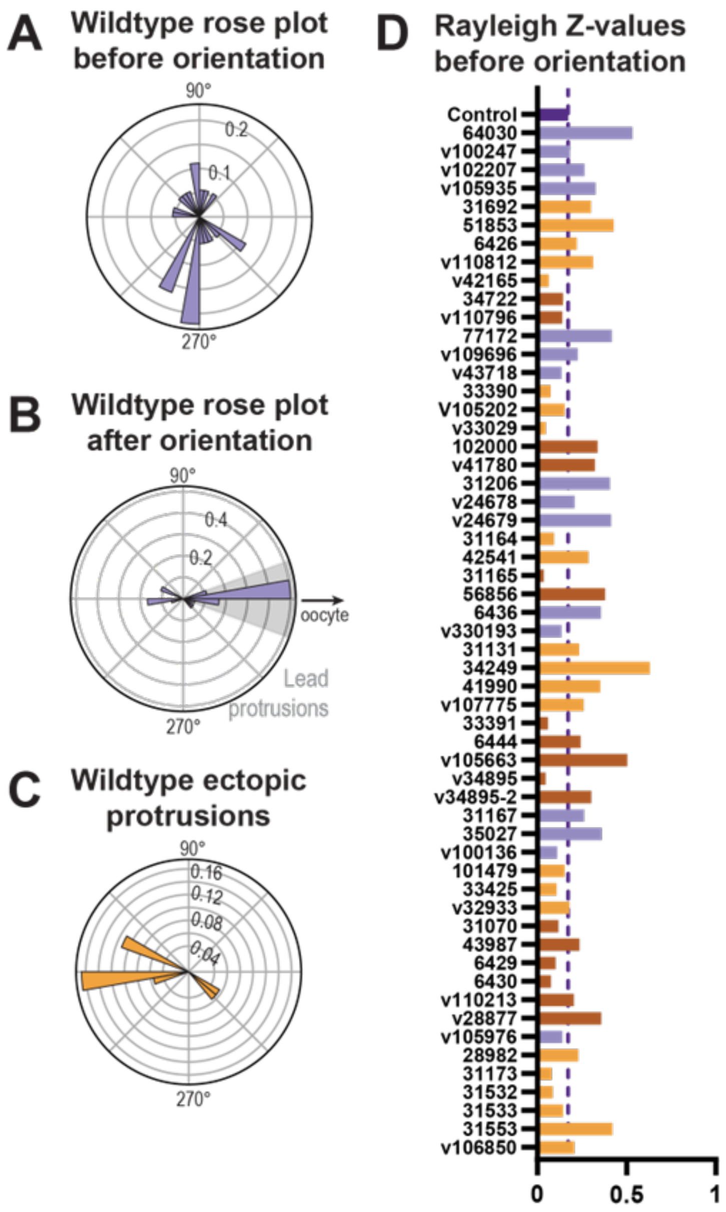


**Supplemental Figure 4: Border cell clusters require orientation prior to analysis to correctly quantify directionality of protrusions. (A)** Aggregated protrusions for wildtype clusters before orientation cannot be used to differentiate lead versus ectopic protrusions. **(B)** Orienting the clusters shows a strong directional preference in wild-type clusters. **(C)** Excluding the lead protrusions allows quantification of ectopic protrusions and their directionality. **(D)** Rayleigh Z-values before orientation are low, demonstrating a lack of directional preference and highlighting the need for orientation.
